# *SCN1A*-deficient hiPSC-derived excitatory neuronal networks display mutation-specific phenotypes

**DOI:** 10.1101/2023.01.11.523598

**Authors:** Eline J.H. van Hugte, Elly I. Lewerissa, Ka Man Wu, Giulia Parodi, Torben van Voorst, Naoki Kogo, Jason M. Keller, Dirk Schubert, Helenius J. Schelhaas, Judith Verhoeven, Marian Majoie, Hans van Bokhoven, Nael Nadif Kasri

**Affiliations:** Department of Human Genetics, Radboudumc, 6500 HB Nijmegen, The Netherlands; Donders Institute for Brain, Cognition, and Behaviour, 6500 HB Nijmegen, The Netherlands; ACE Kempenhaeghe, Department of Epileptology, 5591 VE Heeze, The Netherlands; Department of Informatics, Bioengineering, Robotics, and Systems Engineering (DIBRIS), University of Genova, 16145 GE Genova, Italy; Stichting Epilepsie Instellingen Nederland (SEIN), 2103 SW Heemstede, The Netherlands

**Keywords:** Epilepsy, MEA, hiPSC

## Abstract

Dravet syndrome is a severe epileptic encephalopathy, characterized by (febrile) seizures, behavioral problems and developmental delay. 80% of Dravet syndrome patients have a mutation in *SCN1A*, encoding Na_V_1.1. Milder clinical phenotypes, such as GEFS^+^ (generalized epilepsy with febrile seizures plus), can also arise from *SCN1A* mutation*s*. Predicting the clinical phenotypic outcome based on the type of mutation remains challenging, even when the same mutation is inherited within one family. Both this clinical and genetic heterogeneity add to the difficulties of predicting disease progression and tailored prescription of anti-seizure medication. A better understanding of the neuropathology of different *SCN1A* mutations, might give insight in differentiating the expected clinical phenotype and best fit treatment choice. Initially it was recognized that loss of Na^+^ -current in inhibitory neurons specifically resulted in disinhibition and consequently seizure generation. However, the extent to which excitatory neurons contribute to the pathophysiology is currently debated, and might depend on the patient clinical phenotype or the specific mutation in *SCN1A*.

To examine the genotype-phenotype correlations of *SCN1A* mutations in relation to excitatory neurons, we investigated a panel of patient-derived excitatory neuronal networks differentiated on multi-electrode arrays. We included patients with different clinical phenotypes, harboring different mutations in *SCN1A*, plus a family where the same mutation leads to both GEFS+ and Dravet syndrome.

We hitherto describe a previously unidentified functional excitatory neuronal network phenotype in the context of epilepsy, which corresponded to seizurogenic network prediction patterns elicited by proconvulsive compounds. We find that excitatory neuronal networks were differently affected, dependent on the type of S*CN1A* mutation, but not on clinical severity. Specifically, pore domain mutations could be distinguished from voltage sensing domain mutations. Furthermore, all patients showed aggravated neuronal network responses upon febrile temperatures. While the basal neuronal network phenotypes could not be distinguished based on patient clinical severity, retrospective drug screening revealed that anti-seizure medication only affected GEFS+ patient-, but not Dravet patient-derived neuronal networks in a patient specific and clinically relevant manner. In conclusion, our results indicate a mutation-specific excitatory neuronal network phenotype, which recapitulates the foremost clinically relevant features, providing future opportunities for precision therapies.

**Highlights:**

- Human stem cell derived excitatory neurons are affected by mutations in *SCN1A* and display mutation-specific, but not clinical phenotype specific, neuronal network phenotypes
- The neuronal network phenotype we describe corresponds to seizurogenic network prediction patterns in vitro
- Excitatory neuronal networks respond to Dravet syndrome clinically relevant triggers, like febrile temperatures and Dravet-contraindicated ASM Carbamazepine
- Retrospective drug screening revealed that GEFS+ neuronal networks, but not Dravet neuronal networks respond to ASM in a patient-specific and clinical relevant manner

## Introduction

Dravet syndrome (DS) is a severe epileptic encephalopathy characterized by recurrent and prolonged seizures, manifesting before the first year of life^1^. Patients have a poor prognosis, and the phenotype progresses into complex seizures and neurodevelopmental delay including behavioral, cognitive and motor impairments^1–4^. In 80% of the cases, DS is caused by mutations in *SCN1A*, encoding the voltage gated Na^+^ -channel NaV1.1^5,6^. Also a milder clinical phenotype, generalized epilepsy with febrile seizures plus (GEFS^+^), can arise from *SCN1A* mutations. Most patients with GEFS^+^ show normal development and spontaneously remitting seizures later in life^2^.

To date, more than 1500 different *SCN1A* mutations have been identified, most of which occur *de novo*^7,8^, including missense, frame shifts, splice site variants, or insertions/deletions that lead to loss-of-function (LoF)^9,10^. Although evidence suggests that the nature of the mutation affects the phenotype^9^, with LoF mutations conferring a higher risk ^11^, predictions based on individual mutations remain challenging. The same mutation can give rise to variable phenotypes even if they segregate within one family^12^. Both this clinical and genetic heterogeneity add to the difficulties of predicting disease progression and prescriptions of patient-tailored anti-seizure medication (ASM).

*Scn1a* knock out mouse models for DS showed that a loss of excitability, caused by a reduced Na^+^ -current density, predominantly affect parvalbumin-positive and somatostatin-positive inhibitory neurons^13–18^. Disinhibition is considered the major pathogenic mechanism of DS, in turn leading to hyperexcitability and epileptic activity^19^. However, depending on the strain or age of the mice, excitatory neurons equally contribute to the epileptic activity^20^. Studies using human induced pluripotent stem cells (hiPSC)-derived neurons provided further conflicting results. While some reported deficits in Na^+^ -current density or action potential firing in inhibitory neurons alone^21–24^, others reported deficits in both excitatory and inhibitory neurons^25–27^.

These conflicting observations might be attributed to intrinsic functional differences of the investigated mutations or the cell types studied. Some mutations result in gain-of-function (GoF), while others in (partial) LoF of Na_V_1.1, suggesting that different *SCN1A* variants might affect inhibitory or excitatory neurons to different extend^8^. However, it remains difficult to attribute the neuronal phenotypes either to the patient clinical phenotype, or to the type of variant studied. Moreover, studies have not systematically investigated the effect of individual mutations at the neuronal network level. Only a series of patient hiPSC-derived neurons, that include multiple *SCN1A*-variants and clinical phenotypes, can uncover if the excitatory neuronal phenotype is dependent on the patient clinical phenotype or the type of mutation studied.

To this end, we investigated CRISPR/Cas9-edited *SCN1A* -lines, and a panel of hiPSC-derived lines from five patients with different clinical phenotypes, harboring different *SCN1A* variants, and mapped their effect on excitatory neuronal network behavior. In particular, we studied a family where the same *SCN1A* mutation resulted in different clinical phenotypes. Our data provide evidence that excitatory neurons are affected by different *SCN1A* mutations to different extend. Specifically, missense versus nonsense mutations in *SCN1A*, and the location of the mutation in Na_V_1.1, pore domain versus voltage sensing domain, influences the excitatory network phenotype. Although we did not observe differences based on clinical severity in basal network phenotypes, DS clinical relevant triggers, like febrile temperatures and ASM, elicited different responses in DS and GEFS+ -patient derived neuronal networks. We here provide a mutation-specific disease model, presenting future opportunities for precision therapies in the context of DS

## Methods

### Patient inclusion and hiPSC generation

All experiments involving peripheral mononuclear blood cells (PBMCs) and hiPSCs from DS patients were carried out after informed consent and approval by the medical ethical committee of the Radboudumc, Nijmegen (2018-4525). From all subjects, PBMCs were isolated via a blood sample scheduled during routine diagnostic testing. Patient clinical history is described in the supplementary methods.

PAT001_DRAV was a 4-year-old female at time of sampling. She had a missense mutation (c.4168G>A p.Val1390Met) in the pore domain of *SCN1A*. PAT001_GEFS was a female patient and 10 years at the time of sampling, with a missense mutation (c.2576G>A p.Arg859His) in the voltage sensing domain. FAM001_DRAV, FAM001_GEFS and FAM001_FS all belonged to the same family, and all carried a mutation in c.3926T>G (p. Leu1309Arg) in the voltage sensing domain. FAM001_DRAV was the male sibling of 16 years and FAM001_GEFS was the female sibling of 14 years at time of sampling. FAM001_FS was the father and was 49, CTRL1 was the unaffected mother and was 43 at time of sampling.

All lines were reprogrammed using episomal vectors with the Yamanaka transcription factors *Oct4, c-Myc, Sox2* and *Klf4* ^28^, and had normal karyotypes. All mutations were confirmed via sanger sequencing. CTRL2 was an external control derived from a 30-year-old male, previously characterized^29^, and tested for genetic integrity using SNP assay^30^.

### CRISPR/Cas9 gene editing of *SCN1A*

From CTRL2, two *SCN1A* CRISPR lines were generated (*SCN1A^+/-^* 3, and *SCN1A^+/-^* 7) with a heterozygous deletion in exon *21*, leading to a premature stop codon at amino acid position 18. *SCN1A^+/-^* 3 had a heterozygous mutation at position c.500_501insAC (p.Thr18*) and *SCN1A^+/-^* 7 at position c.500_501delGTCAinsTACTC (p.Thr18*). A short guide RNA (sgRNA) was designed (GTCAAACCTGTCACCAGTTG) and cloned into pSpCas9(BB)-2A-Puro (PX459) V2.0 (Addgene, #62988) according to a previously published protocol ^31^. In brief, 8 x 10^5^ hiPSCs in single-cell suspension were nucleofected with 5 μg SpCas9-sgRNA plasmid using the P3 Primary Cell 4D-Nucleofector Kit (Lonza, #V4XP-3024) in combination with the 4D Nucleofector Unit X (Lonza, #AAF-1002X). Cells were resuspended in E8 Flex supplemented with Revitacell (10 μg/ml, Thermo Fisher, #A2644501).) and seeded on Biolaminin 521 LN (20 μg/ml, BioLamina, #LN521-05) pre-coated wells. 24 hours after nucleofection, 0.5 μg puromycin (Sigma-Aldrich, #P9620) was added for 24h. Resistant colonies were manually picked and send for Sanger sequencing to ensure heterozygous editing of exon 21.

### hiPSC culturing and neuronal differentiation

All hiPSC lines were cultured on Matrigel (Corning, #356239) in Essential 8 (E8) flex medium (Gibco, #A2858401) supplemented with primocin (0.1 μg/ml, Invivogen, #ant-pm-2). For quality check, hiPSCs were tested for pluripotency using qPCR or flow cytometry (Fig S1K, L). hiPSCs were infected according to a previously published protocol with lentiviral constructs encoding *rtTA* combined with *Ngn2* to generate doxycycline-inducible excitatory neurons^32,33^. Selection was performed by adjusting the concentrations of G418 (100 μg/ml - 250 μg/ml, Sigma-Aldrich, #G8168), for *Ngn2*, and puromycin (1 μg/ml - 2 μg/ml), for rtTA. Cells were passaged twice a week using ReLeSR passaging agent (Stemcell, #05872), and not kept for more than 10 passages. Every 2 weeks, cells were checked for mycoplasma contamination.

hiPSCs were differentiated to glutamatergic neurons by overexpression of *Ngn2* and cultured in a 1:1 ratio with E18 rodent astrocytes according to a previously published protocol^32,33^. Detailed neuronal differentiation protocol can be found in the supplementary methods.

### Immunocytochemistry

Coverslips were washed once with ice-cold PBS. Unless stated otherwise, subsequent steps are performed at room temperature and solutions were made in PBS. Cells were fixated with 4% PFA and 4% sucrose for 15 minutes. Fixed coverslips were washed tree times, and stored in PBS at 4°C until further use. Cells were permeabilized with triton X-100 (0.2%, Sigma-Aldrich, #T8787) for 10 minutes, followed by 1 hour of blocking with normal goat serum (NGS) (5%, Invitrogen, #10000C). Primary antibodies raised against Na_V_1.1 (1:500, Alomone labs, #ASC-001) MAP2 (1:1000, Synaptic Systems, #188004) and AnkG (1:1000, Invitrogen, #33-800) diluted in 1% NGS were incubated overnight at 4°C. After 10 washes with PBS, coverslips were incubated in secondary antibodies diluted in 1% NGS. Coverslips were washed 10 times with PBS, and stained with Hoechst (0.01%, ThermoFisher, #H3570). The coverslips were mounted in DAKO (Agilent, #S3023) on microscope slides, and imaged on a Zeiss Axio Imager Z2 at 63x magnification.

### Western Blot

For Western Blot, a total of 350.000 cells were seeded on a 6 well plate. At day in vitro (DIV) 49, proteins were extracted in cold RIPA lysis buffer and supernatant was used to determine protein concentration using the Pierce BCA protein assay kit (Thermofisher). 50 μg protein was loaded on Mini-PROTEAN TGX Stain-Free Gels before transfer to Nitrocellulose membranes (Biorad). The membrane was blocked with blotto blocking buffer (Santa cruz, #sc-2333) for 1 hour at room temperature and incubated with primary antibody overnight at 4°C against Na_V_1.1 (1:500), using GAPDH (1:1000, CellSignaling, #2118S) as a loading control. Membranes were washed three times in TBS-T followed by incubation for 1 hour with Goat anti-Rabbit IgG (H+L) secondary antibody, HRP (1:50.000, Thermo Fisher, #G-21234) at room temperature. Membranes were washed five times in TBS-T, before ECL detection with the Supersignal West Femto Kit (Thermo Fisher, #34095) and imaged on the Biorad Gel Imaging system (Chemidoc) and subsequently analyzed in ImageJ ^34^.

### MEA recording and analysis

To record spontaneous network activity, multiwell-MEAs were used that consisted of 24 individual wells (Multichannel systems, MCS GmbH, Reutlingen, Germany) according to previous published protocol ^35^. In brief, each MEA well was embedded with 12 gold electrodes spaced 300 μm apart with a diameter of 30 μm. Baseline neuronal network activity was recorded for 10 minutes at DIV 49, after a 10 minute acclimatization period, in a recording chamber maintained constant at 37°C/95% O_2_/5% CO_2_. For temperature experiments, the amplifier temperature was set at 40°C. When this temperature was reached, the cultures were allowed to acclimate for 10 minutes before a 10 minute recording at 40°C. The same method was applied for the recovery recording at 37°C. For all recordings, the raw signal was sampled at 10 kHz and filtered with a high-pass 2^nd^ order Butterworth filter with a 100 Hz cut-off and a low-pass 4^th^ order Butterworth filter with a 3500 Hz cut-off. The noise threshold for individual spike detection was set at ±4.5 standard deviations.

*Pharmacology* All reagents were prepared into stocks and stored at −20°C. A set of 4 proconvulsive compounds were chosen to induce seizurogenic responses of the neuronal network at DIV 49: NMDA (100 μM working concentration, Tocris, #6384-92-5), Linopirdine (1.5 μM, Sigma, #105431-72-9), 4-AP (10 μM, Tocris, #504-24-5), Phenytoin (25 μM, Sigma, #57-41-0) and Kainic acid (10 μM, Sigma #58002-62-3). The working dilution was directly added to the wells during MEA recordings. A set of 4 ASM was used to rescue neuronal network phenotypes. Cultures were treated from DIV13 onwards, and ASM concentration was kept constant at 10 μM: Valproic acid (Tocris, #1069-66-5), Levetiracetam (Tocris, #102767-28-2), Topiramate (Tocris, #97240-79-4), and Carbamazepine (Tocris, #298-46-4). The working dilution was first diluted into the medium before regular medium change. For all pharmacological dilutions, the amount of DMSO in the cell culture medium was ≤0.5% v/v in each experiment.

#### Data analysis using Multiwell-Analyzer

To guarantee sufficient experimental quality, we adhered to a set of published guidelines ^35^. We included experiments with a minimum of 9 wells per hiPSC line, across at least two independent batches. Networks that did not show network bursts (NBs) at DIV 27, or MFR <0.1 Hz and burst rate <0.4 bursts per minute were excluded. Wells that displayed low cell densities or cell clumping, were discarded. All experiments were carried out at DIV 49. Off-line data analysis was performed using Multiwell-Analyzer software (Multichannel systems) that permitted the extraction of spike-trains, and a custom-made in-house code developed in MATLAB (The Mathworks, Natick, MA, USA). The mean firing rate (MFR) (Hz) was calculated for each well individually by averaging the firing rate of each separate channel by the total amount of active channels of the well. The Multiwell analyzer build-in burst detection algorithm was used to detect bursts, and defined bursts if 4 spikes were in close proximity with a maximum of 50 ms inter spike interval (ISI) to start a burst, a maximum of 50 ms ISI to end a burst, and a minimum of 100 ms inter burst interval (IBI). NBs were defined when a burst was present in at least 50% of all channels. The percentage of random spikes (PRS) was defined by calculating the percentage of isolated spikes that did not belong to a burst. The IBI, and NB IBI (NIBI) were calculated by subtraction of the time stamp of the beginning from the time stamp of the ending of each burst or NB. The mean IBI per channel was calculated and averaged across all channels. The NIBI was averaged for each well. High frequency bursts (HFB) inside a network bust period were detected by decreasing the burst detection ISI to 5 ms, with a maximum IBI of 10 ms, to individually detect each HFB. An in-house Matlab script quantified the number of HFBs inside one network bust period using a maximum inter-HFB-interval of 1500 ms.

### Single cell electrophysiology

Experiments were performed in a recording chamber at DIV49, and visualized using an Olympus BX51WI upright microscope (Olympus Life Science, PA, USA) and a DAGE-MTI IR-1000E (DAGE-MTI, IN, USA) camera. Dara were acquired using a Digidata 1440-A digitizer and a Multiclamp 700B amplifier (Molecular Devices, San Jose, CA, USA), sampled at 20.000 Hz, and filtered using a low-pass 1 kHz filter. Filamented patch pipettes were pulled from borosilicate glass (Science Products GmbH, Hofheim, Germany) with a PC-10 micropipette puller (Narishige, London, UK), with an open tip resistance between 5-7 MΩ. The recording chamber was continuously perfused with artificial cerebrospinal fluid (ACSF), containing (in mM): 124 NaCl, 1.25 NaH2PO4, 3 KCl, 26 NaHCO3, 11 Glucose, 2 CaCl2, 1 MgCl2 (adjusted to pH 7.4), and maintained constant at 37°C/5% CO_2_. Recordings were excluded if series resistance was above 20MΩ, or if the series resistance to membrane resistance was below a 1:10 ratio.

#### Intrinsic properties

For intrinsic properties, pipettes contained a potassium-based internal solution containing (in mM): 130 K-Gluconate, 5 KCl, 10 HEPES, 2.5 MgCl2, 2 Na2-ATP, 0.4 Na3-ATP, 10 Na-phosphocreatine, 0.6 EGTA (adjusted to pH 7.25 and osmolarity 290 mOsmol). Resting membrane potential (vRMP) was determined immediately after reaching whole cell configuration in current clamp. Both active and passive membrane properties were determined at a holding potential of −60 mV. Action potentials were elicited by a stepwise current injection protocol ranging from −30pA to +50pA and determined by analyzing the first action potential that was elicited. All intrinsic properties were analyzed using Clampfit 10.7 (molecular devices).

#### Na^+^ -currents

A CsCl based internal pipette solution was used, containing (in mM): 115 CsMeSO_3_, 20 CsCl, 10 HEPES, 2.5 MgCl_2_, 4 Na_2_ATP, 0,4 Na_3_GTP, 10 Na-phosphocreatine, 0.6 EGTA (pH adjusted to 7.24, osmolarity 294 mOsmol). CNQX (40 μM, Tocris #0190) was used to block synaptic activity during recording. TTX (1 μM, Tocris, #1069) was used as a negative control. Cells were measured in whole cell voltage clamp configuration including P/8 leak subtraction. Using a step wise protocol, cell membrane potential was increased from a holding potential of −90 mV in 10 mV increments to 60 mV for 100 ms. Peak currents were determined using Clampfit 10.7, and divided by the cell capacitance. Conductance was calculated using the following equation: G = I/(V-V_rev_), where G is the conductance, I the peak current amplitude, V the holding potential of each pulse, and V_rev_ the Na^+^ reversal potential. V_rev_ was extrapolated from the experimental data using an in house Matlab script, and data was fitted to the following Boltzmann equation: I = Imax/(1+exp((V-V_1/2_)/*k*)), where I is the normalized current, Imax the maximum current, V the holding potential of each pulse, V1/2 the half maximum of activation and *k* the slope.

### Statistics

Statistical analysis was performed using Graphpad PRISM 9.0.0 (GraphPad Software, Inc., CA, USA). All values are reported as mean ± SEM, unless stated otherwise. In all figures, *P*-values are indicated as follows: <0.05 (*), <0.005 (**), <0.0005 (***), <0.0001 (****), two-tailed. Bonferroni correction for multiple testing was performed when needed. Principal component analysis (PCA) was performed on data from 8 parameters (MFR, PRS, BR, BD, BSR, NBR, NBD, HFB) using Graphpad PRISM 9.0.0. Values were standardized and principal components were selected based on parallel analysis using Monte Carlo simulation on random data of equal dimension to the input data, generated with a random seed.

## Results

### Proconvulsive compounds induce three distinct neuronal network phenotypes in control neuronal networks

Previously, we functionally benchmarked ten control neuronal networks on multi-electrode arrays (MEAs), by differentiating hiPSCs to electrophysiologically mature excitatory neurons through forced expression of *Ngn2*^32,33,35^. Network activity consisted of spikes (single action potentials) and bursts (clustered action potentials at high frequency), which self-organized into synchronous NBs (bursts recorded from all electrodes) (Fig. 1A)^35^. Traditionally, proconvulsive compounds are used to elicit neuronal network responses as a predictor for seizurogenic network activity^36,37^. To understand the applicability of excitatory neurons as a model for DS-related epilepsy, and to predict excitatory seizurogenic phenotypes, we selected five different seizurogenic compounds that affect excitatory neuronal mechanisms and functionally characterized the network properties (Fig. 1B-J). N-methyl-D-aspartate (NMDA) and Kainic acid (KA), both led to a hyperactive neuronal network phenotype, reflected by a significant increase in mean firing rate (MFR), burst rate (BR) and NB rate (NBR) (Fig. 1B-I). K^+^ -channel blockers 4-AP and Linopirdine, showed a distinct seizure prediction pattern (Fig 1B, J). While the neuronal networks responded with a similar increase in MFR (Fig. 1C), the organization of the network differed, and showed an irregular neuronal network pattern (Fig. 1B). Whereas the NBR was not significantly increased, the NB duration (NBD) increased, and the NBs consisted of several high frequency bursts (HFB) (Fig 1E-I). Na^+^-channel blocker phenytoin, led to a third distinct seizure prediction pattern, with a general decrease in activity and synchronicity, reflected by a decreased MFR, BR and NBR, but an increased percentage of random spikes (PRS, isolated, a-synchronous spikes) (Fig 1C-E, H). A heatmap clearly showed three different seizure prediction patterns upon exposure to proconvulsive compounds (Fig. 1J), in analogy to neuronal networks in rodent cultures^37^. Prediction pattern 1, elicited by KA and NMDA lead to a hyperactive neuronal network phenotype. Prediction pattern 2, elicited by phenytoin led to a hypoactive, desynchronized network activity. Finally, prediction pattern 3, elicited by 4-AP and Linopirdine led to disorganization including NBs with HFB.

**Figure 1.**
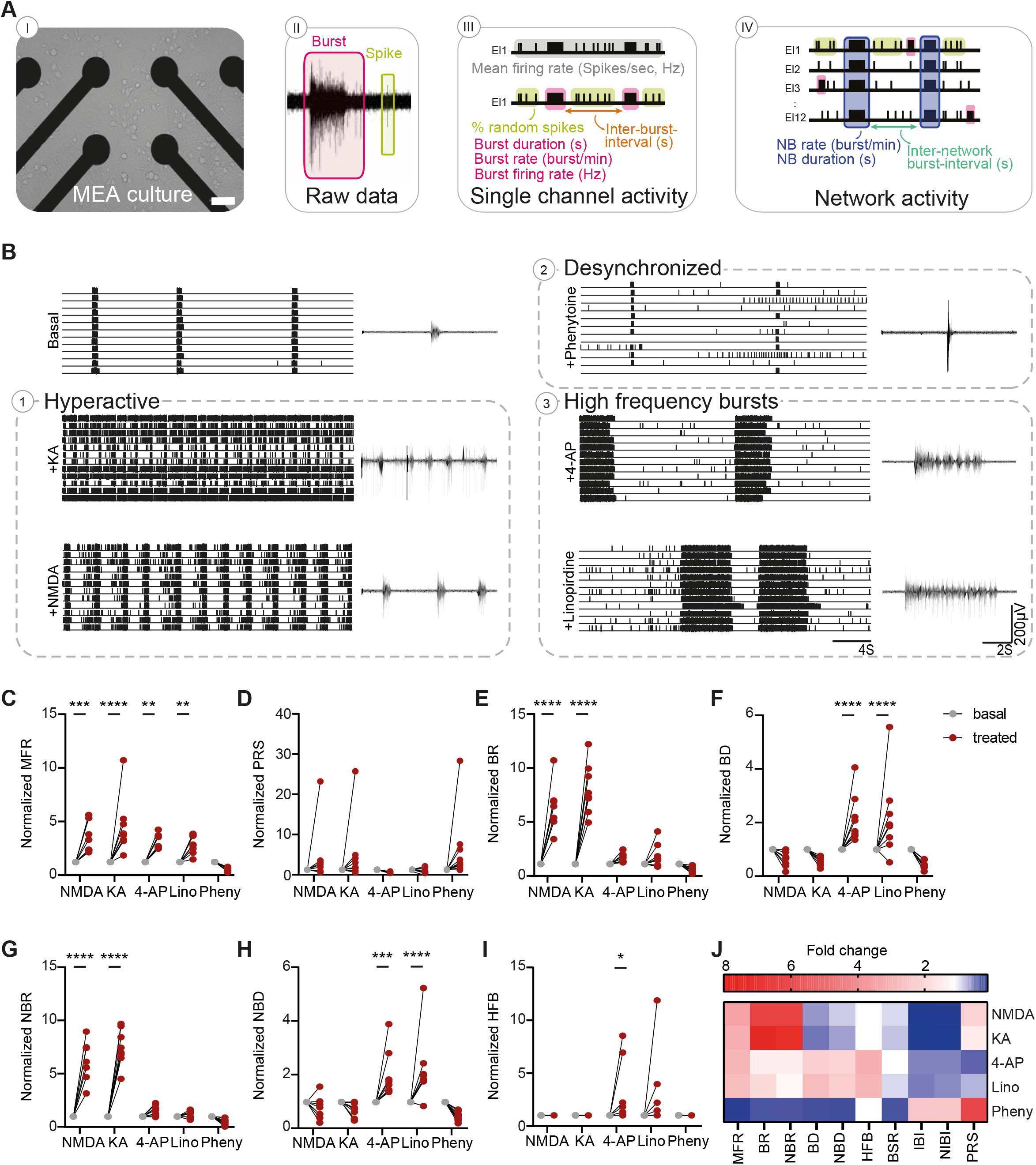
Seizure liability screening reveals two distinctive seizure prediction patterns. A) Schematic overview of a MEA culture, raw recording from a representative electrode and rasterplot including the measured MEA parameters on both single channel and network levels. Scale bar represents 100 μm B) Representative rasterplots of MEA recordings from control networks and control networks acutely treated with different compounds, NMDA (N-methyl-D-aspartate) (100 μM), 4-AP (4-Aminopyridine) (10 μM), KA (Kainic acid) (10 μM), Lino (Linopirdine) (1.5 μM), and Pheny (Phenytoin) (25 μM) with 8s zoom-in of the activity of one electrode. Quantification of network parameters normalized to pre-treatment baseline including: C) mean firing rate (MFR), D) percentage of random spikes (PRS), E) burst rate (BR), F) burst duration (BD), G) network burst rate (NBR), H) network burst duration (NBD), and I) number of high frequency bursts (HFB). J) Heatmap of all MEA parameters for all compound-treated recordings, normalized to pre-treatment baseline, including MFR, BR, NBR, PRS, BD, NBD, HFB, burst spike rate (BSR), interburst interval (IBI) and network inter burst interval (NIBI). For all MEA data NMDA n=7, KA n=8, 4-AP n=8, Lino n=8 and Pheny n=10 (pre- and posttreatment wells), \**P* = 0.05, \*\**P* = 0.01, \*\*\**P* = 0.001, \*\*\*\**P*<0.0001, Two-way ANOVA with Dunn–Šidák multiple comparisons correction. All means, SEM and test statistics are listed in table 1.

### Heterozygous LoF mutations in *SCN1A* lead to de-synchronized and hyperactive neuronal networks in excitatory neurons

Heterozygous LoF mutations in *SCN1A* predominantly result in a severe DS clinical phenotype, and not in an intermediate GEFS^+^- or mild febrile seizure (FS)- like phenotype^11^. To examine the effect of heterozygous LoF of *SCN1A* in mature excitatory neuronal networks, we generated two *SCN1A^+/-^* - deficient lines (*SCN1A*^+/-^ Cl3, and *SCN1A*^+/-^ Cl7). We used CRISPR/Cas9 genome editing in a curated control hiPSC-line to induce a frameshift mutation, leading to a heterozygous deletion in exon 21 of *SCN1A*, resulting in a premature stop codon at amino acid position 18 in both clones (Fig. S1A-C). Immunocytochemistry confirmed the expression of Na_V_1.1 in excitatory neurons (Fig. S1D). In line with the localization of Na_V_1.1 in inhibitory neurons, we found Na_V_1.1 expression in the soma and axon initial segment, with specific dense puncta^16^ (Fig. S1DII). In addition, Na_V_1.1 was expressed in the dendrites (Figure S1DI). Western blot confirmed reduced Na_V_1.1 expression in *SCN1A^+/-^* -deficient lines (Fig. S1E).

Control networks showed synchronous, rhythmic NBs at day DIV49 (Fig. 2A), a hallmark of physiological network activity^35^. In contrast, *SCN1A^+/-^* -deficient lines showed a hyperactive, de-synchronous network state (Fig. 2A), where NB regularity was completely disrupted. The network activity in both *SCN1A^+/-^* -deficient lines showed blocks of intense high-frequency periods, consisting of multiple HFB, followed by relatively silent periods, and a significant increase in PRS, MFR, NBR and BR (Fig. 2B, Fig. S1F), but a significantly decreased (N)BD (Fig. S1F). Principal component analysis (PCA) showed clear separation of *SCN1A^+/-^* -deficient and control networks (Fig. 2C). In conclusion, *SCN1A^+/-^* -deficient neuronal networks were differently organized than control neuronal networks, showing a hyperactive-desynchronized neuronal network phenotype, resembling previously identified seizure prediction pattern 1 (Fig 1J).

**Figure 2.**
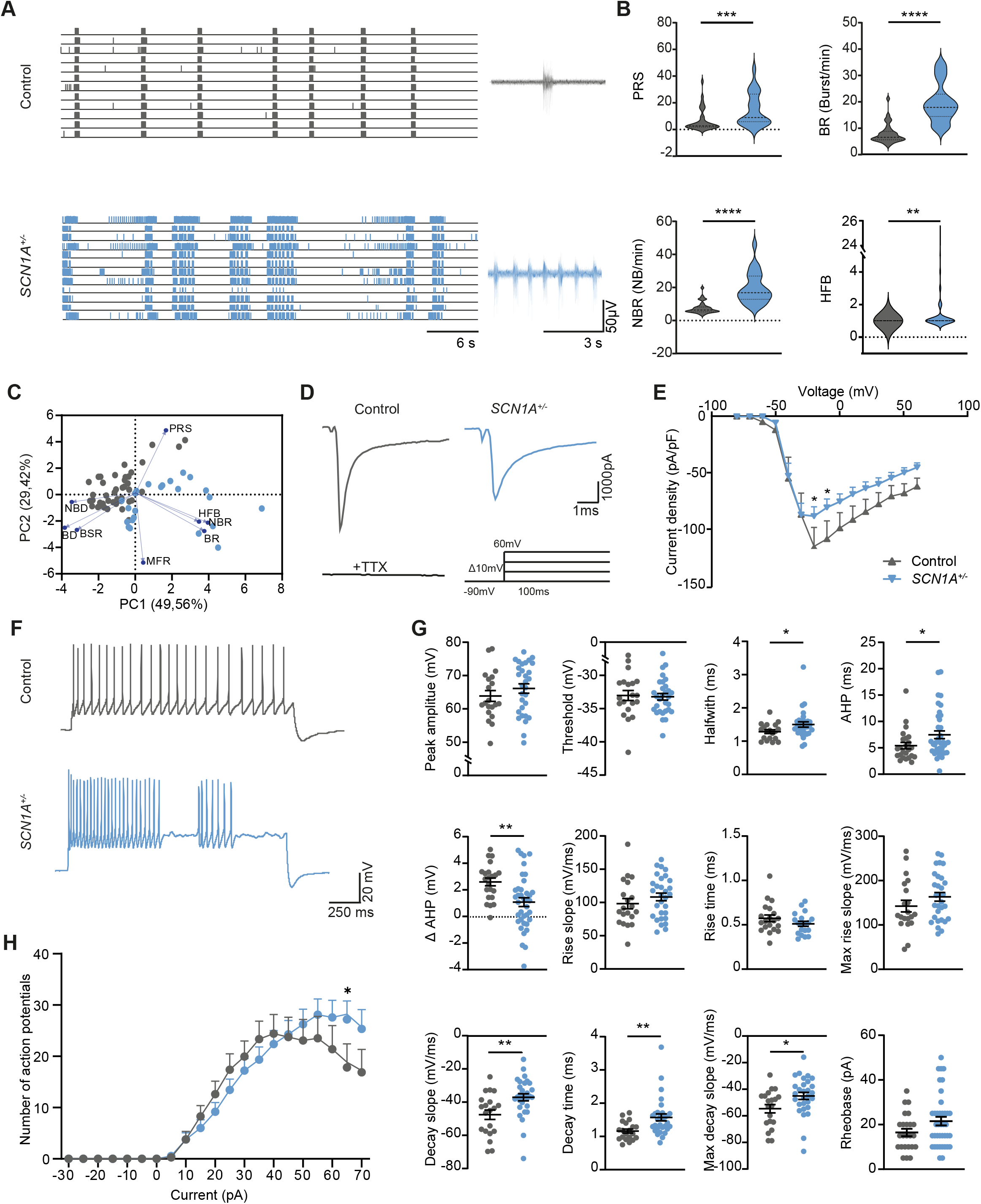
*SCN1A^+/-^* -deficient neurons depict hyperactive, de-synchronized and irregular firing patterns. A) Representative rasterplots of MEA recordings at DIV49 from control and *SCN1A^+/-^* -deficient neuronal networks, with 6s zoom-in of the activity of one electrode. B) Quantification of network parameters including: percentage of random spikes (PRS), burst rate (BR), network burst rate (NBR), and number of high frequency bursts (HFB), quantified as the number of HFBs inside a network burst period. Dashed line represents median, dotted line represents quartiles. C) PCA-plot of 8 MEA parameters, including mean firing rate (MFR), BR, NBR, PRS, burst duration (BD), NBD, HFB, burst spike rate (BSR), showing parameters that explain the differences in network behavior between control and *SCN1A^+/-^* -deficient neuronal networks. Blue arrows indicate loadings. For all MEA data, n = number of recordings/batches: control n = 33/5 and *SCN1A^+/-^* n = 23/3, Mann-Withney test. D) Representative Na^+^ -current traces of control and *SCN1A^+/-^* -deficient neurons at DIV49. Stimulation paradigm (inset): stepwise protocol from a −90 mV holding potential to a maximum test-pulse of 60 mV in increments of 10 mV, Na^+^ -currents were Tetrodotoxin (TTX) sensitive. E) Current-density plot of Na^+^ -current recordings from n = cells/batches Control n = 19/3 and *SCN1A^+/-^* n = 22/3 neurons, 2-way ANOVA with Dunett’s multiple comparisons test. F) Representative firing patterns of control and *SCN1A^+/-^* neurons, measured by a step-wise current injection protocol. G) Analysis of active and passive membrane properties including: peak amplitude, threshold, halfwidth, after hyper polarization time (AHP), ΔAHP, defined as the difference between the AHP of the first action potential and the second action potential in the same sweep, rise slope, rise time, maximum rise slope, decay slope, decay time, max decay slope, rheobase. H) quantification of the number of action potentials per current injection. For all action potential properties, n = cells/batches control n = 22/3 and *SCN1A^+/-^* n = 30/3, Mann-Withney test. Data represented as mean ±SEM. \**P* = 0.05, \*\**P* = 0.01, \*\*\**P* = 0.001, \*\*\*\**P*<0.0001. All means, SEM and test statistics are listed in table 1.

To determine the contribution of single cell properties to the phenotype observed in *SCN1A^+/-^* -deficient excitatory networks, we determined Na^+^-current and action potential properties (Fig. 2D-H, Fig. S1G-J). Current/voltage plots showed a slight reduction in Na^+^ -current density in *SCN1A^+/-^* - deficient neurons (Fig. 2D,E), without changes in the channel kinetics (Boltzmann-fitting) (Fig S1H). Action potentials in *SCN1A^+/-^* -deficient neurons had a significantly increased halfwidth (Fig. 2G), which is directly related to fast neuronal Na^+^ -channel activation. Interestingly, this was not reflected by altered rise dynamics (Fig. 2G). Rather, action potentials in *SCN1A^+/-^* -deficient neurons showed a significant decline in decay dynamics (Fig. 2G), indicative of altered dynamics in voltage dependent potassium (K_v_) channels. Indeed, the after hyper polarization (AHP) time was increased and inter-spike AHPs (ΔAHP), reflecting AHP adaptation, were more hyperpolarized (Fig. 2G). In addition, *SCN1A^+/-^* - deficient neurons displayed remarkable irregular firing patterns, with a slight increased frequency (Fig. 2F, H, Fig S1J). To conclude, excitatory neurons are affected by *SCN1A^+/-^* -deficiency, both on a neuronal network as a single cell level.

### Patient-specific missense mutations in *SCN1A* lead to a hypo-active and de-synchronized excitatory neuronal network, independent of the clinical phenotype

Previous evidence in both hiPSC-derived neurons and mouse models has shown conflicting data regarding the contribution of excitatory neurons to the DS phenotype^14,16,20,21,26,27^. We here wondered whether the excitatory neuronal network phenotypes segregate according to the type of mutation, and/or the patient clinical phenotype. To this end, we investigated the role of *SCN1A* missense mutations in a patient cohort that consisted of patients with severe (PAT001_DRAV, FAM001_DRAV), intermediate (PAT001_GEFS, FAM001_GEFS) and mild (FAM001_FS) phenotypes (Fig. 3A,B). Particularly, we studied a family where the same missense mutation in *SCN1A* resulted in different clinical phenotypes (FAM001_FS, FAM001_GEFS and FAM001_DRAV) (Fig. 3A,B).

**Figure 3.**
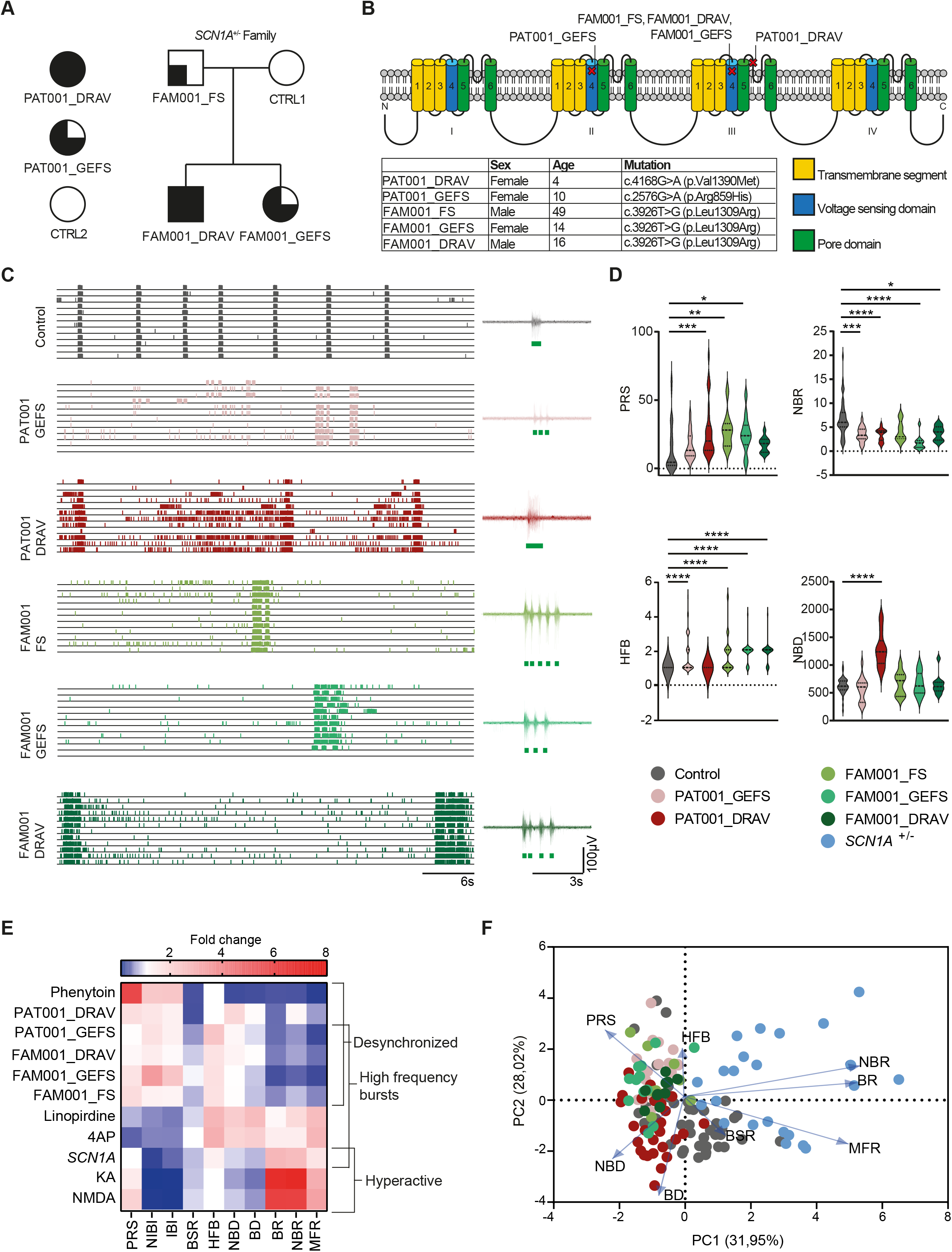
DS patient-derived neuronal networks show mutation specific network fingerprints. A) Schematic overview of patient inclusion, including pedigree of one family, where the same *SCN1A* mutation is inherited via the paternal line. Full black symbol represents DS, three quarters black symbol represents GEFS+ syndrome, one quarter black symbol represents febrile seizures (FS). B) Overview of affected locations in the Na_V_1.1 protein, including a table with patient information and variant position and amino acid changes. C) Representative rasterplots of MEA recordings from control and patient-derived networks, with 6s zoom-in (insets) of the activity of one electrode. D) Quantification of network parameters including: percentage of random spikes (PRS), network burst rate (NBR), number of high frequency bursts (HFB) quantified as the number of HFBs inside a network burst period, and network burst duration (NBD). Dashed line represents median, dotted line represents quartiles. E) Heatmap of all MEA parameters for all compound-treated recordings, normalized to pre-treatment baseline, and all *SCN1A* -deficient lines normalized to control. Parameters include mean firing rate (MFR), burst rate (BR), NBR, PRS, burst duration (BD), NBD, HFB, burst spike rate (BSR), interburst interval (IBI) and network inter burst interval (NIBI). F) PCA-plot of 8 MEA parameters, including MFR, BR, NBR, PRS, BD, NBD, BSR and HFB, showing parameters that explain the differences in network behavior between control and patient lines. Blue arrows indicate loadings. For all MEA data, n = number of recordings/batches: CTRL1 n = 36/5, CTRL2 n = 9/2, PAT001_GEFS n = 22/4, PAT001_DRAV n = 33/6 FAM001_FS n = 9/2, FAM001_GEFS n = 11/3, FAM001_DRAV n = 11/3, *SCN1A^+/-^* n = 23/3., one way ANOVA with Kruskal-Wallis test with Dunss correction for multiple comparisons. \**P* = 0.05, \*\**P* = 0.01, \*\*\**P* = 0.001, \*\*\*\**P*<0.0001. All means, SEM and test statistics are listed in table 1.

All patient-derived lines showed significantly different network phenotypes than control (Fig. 3C). PAT001_DRAV, PAT001_GEFS, FAM001_FS, FAM001_DRAV and FAM001_GEFS neuronal networks consisted of significantly less synchronous NBs and more random spikes (Fig. 3D) than control networks, consistent with seizure prediction pattern 2 (Fig. 1J). Moreover, we observed in PAT001_GEFS, FAM001_FS, FAM001_GEFS and FAM001_DRAV a burst phenotype that included NBs with HFBs, followed by relatively silent periods, reminiscent of the burst phenotype of *SCN1A^+/-^* - deficient lines, and seizure prediction pattern 3 (Fig. 3E). In contrast, PAT001_DRAV neuronal networks did not show this particular HFB phenotype, but presented with a significant increase in NBD (Fig. 3D). Overall, two different network phenotypes were distinguished. We found either a desynchronized neuronal network phenotype, specifically in patient-derived neuronal networks with missense mutations in *SCN1A*, or a de-synchronized but hyperactive neuronal network phenotype, in neuronal networks with a heterozygous LOF in *SCN1A*. The segregation of these different network phenotypes is in line with previously reported seizure-prediction patterns^37^, and the response of the control neuronal network to proconvulsive compounds in figure 1 (Fig. 3E). PCA further revealed four clearly identifiable clusters based on control, *SCN1A^+/-^* -deficient lines and patient lines (Fig. 3F). More importantly, the patient-lines subclustered into patients with a missense mutation in the pore domain of Na_V_1.1 (PAT001_DRAV) and patients with a missense mutation in the voltage sensing domain of Na_V_1.1 (PAT001_GEFS, FAM001_FS, FAM001_GEFS and FAM001_DRAV) (Fig. 3F). We conclude that the excitatory neuronal network phenotype is dependent on the type of *SCN1A* mutation, but that different clinical phenotypes could not be distinguished.

In addition, we investigated if we could uncover mutation-specific phenotypes based on single cell properties. In line with the biophysical properties of a mutation in the pore domain, PAT001_DRAV neurons displayed a significant reduction in Na^+^ -current density (Fig. S2A, B). On the other hand, PAT001_GEFS, FAM001_FS, FAM001_GEFS and FAM001_DRAV excitatory neurons did not show a significant different Na^+^ -current dynamics, V_1/2_ or *k* (S2B, C). Interestingly, the ΔAHP was significantly more hyperpolarized in all patient-derived neurons with a missense mutation in the voltage sensing domain (Fig. S2E).

### Febrile temperatures alter neuronal network organization

As all patients described in this study experienced febrile seizures, we examined the response of the neuronal networks to febrile temperatures (Fig. 4A). Increasing the temperature to 40°C lead to an increase in NBR in control lines, which fully returned to baseline when the temperature was decreased to 37°C (Fig. 4 B, C). Strikingly, all patient-lines that displayed NBs with HFB, showed a significant increase in the number of HFB upon febrile temperatures (Fig. 4D,E). In basal conditions, *SCN1A* family lines could not be distinguished, but after febrile temperature induction only FAM001_FS and FAM001_GEFS neuronal network organization returned to baseline, while FAM001_DRAV neuronal networks did not (Fig 4E). Interestingly, PAT001_DRAV showed a significant higher increase in NBR than the control and other patient networks, signifying a higher sensitivity to febrile temperatures (Fig. 4G), which aggravated during development (Fig. 4H). Moreover, networks measured at febrile temperatures showed more distinguished clusters on PCA based on mutation type (Fig. S2F) than at basal temperatures (Fig. 3F). The response of the patient neuronal network to febrile temperatures resembles the earlier described response of the control network to proconvulsive compounds, with an increase in NBR and HFB as most prominent features, corresponding to prediction pattern 1 and 3, respectively (Fig. 1). To conclude, febrile temperatures alter neuronal network organization, and aggravate the network phenotype associated with HFB.

**Figure 4.**
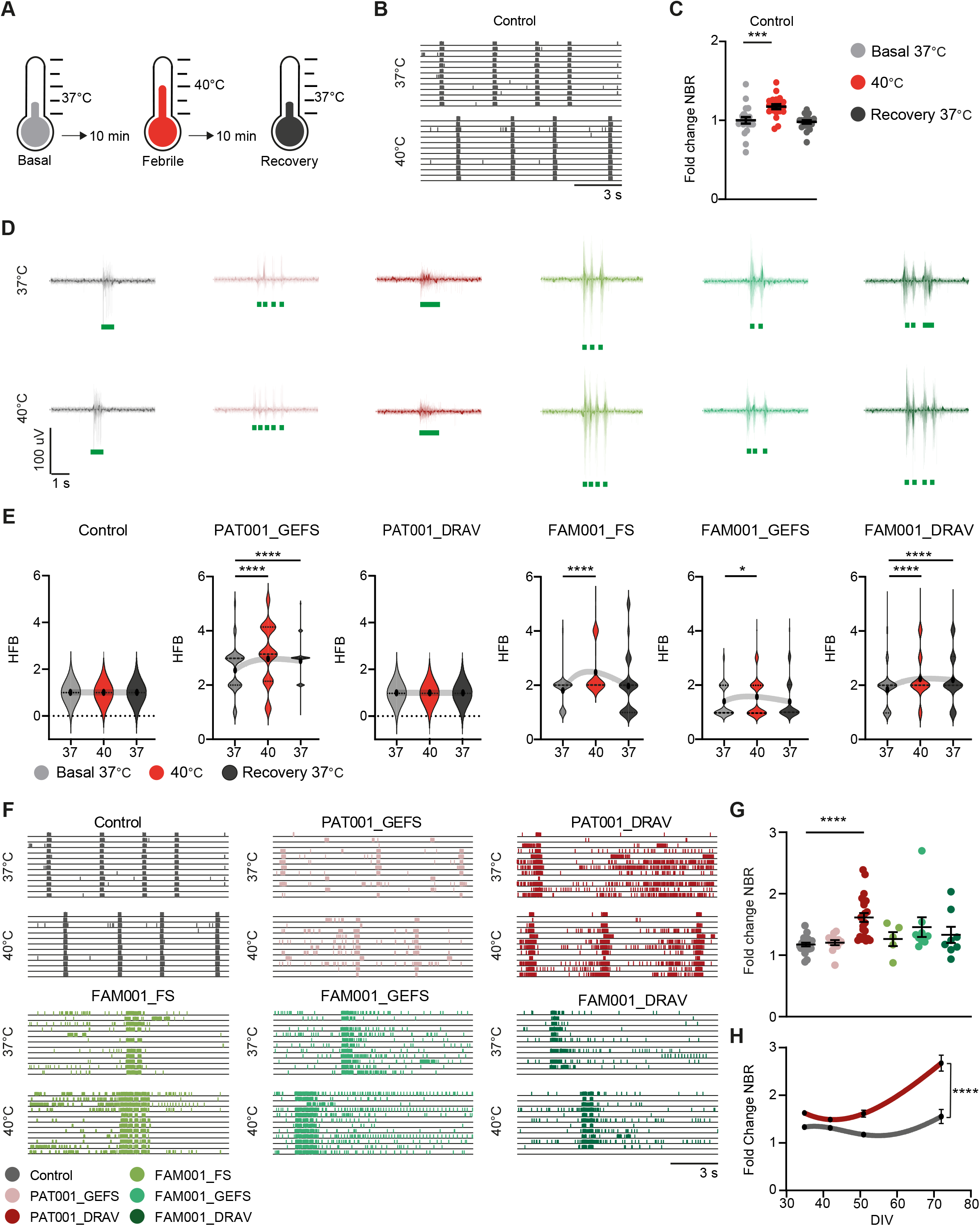
Febrile temperatures lead to altered neuronal network organization. A) Schematic stimulation paradigm for febrile temperature recordings. B) Representative rasterplots from MEA recordings of control neuronal networks, recorded at 37 °C and 40 °C. C) network burst rate (NBR) from control neuronal networks, normalized to the basal recording at 37 °C. Data is shown as mean ±SEM. D) representative burst shapes of control and patient lines green lines indicate high frequency burst (HFB) detection. E) number of HFB, quantified as the number of HFBs inside a network burst period, during basal (37 °C), febrile (40 °C) and recovery (37 °C) recordings. Dashed line represents median, dotted line represents quartiles F) Representative rasterplots from MEA recordings of control and patient neuronal networks, recorded at 37 °C and 40 °C. G) NBR from control and patient neuronal networks, normalized to the basal recording at 37 °C. Data is shown as mean ±SEM. H) NBR from control and PAT001_DRAV neuronal networks over developmental days *in vitro* (DIV), normalized to the basal recording at 37 °C, D significance calculated with two way ANOVA. For all MEA data, N = number of recordings/batches control n = 20/3, PAT001_GEFS n = 12/2, PAT001_DRAV n = 22/3 FAM001_FS n = 5/1, FAM001_GEFS n = 9/2, FAM001_DRAV n = 8/2, all data was recorded at DIV49. \**P* = 0.05, \*\**P* = 0.01, \*\*\**P* = 0.001, \*\*\*\**P*<0.0001, for panel C and G repeated measures ANOVA with Friedmans correction for multiple comparisons was used, for panel E, ANOVA with Kruskal-Wallis test with Dunss correction for multiple comparisons was used. All means, SEM and test statistics are listed in table 1.

### ASM affects GEFS+ neuronal networks in a clinical relevant matter

Although ASM prescribed for DS mostly works on the GABAergic system, we investigated if ASM could also affect the excitatory network phenotype we described, and if excitatory networks already give an indicative prediction of ASM non-responders versus responders. We therefore tested the response of the excitatory neuronal networks to ASM Valproic acid, Levetiracetam, and Topiramate, which all have been prescribed to our patient cohort, and Carbamazepine, which is a general contra-indicated ASM for DS (see supplementary methods). Since HFB was the main phenotypic driver in PAT001_GEFS, FAM001_GEFS and FAM001_DRAV, we compared the effect of the ASM on this parameter, as well as on the NBD in PAT001_DRAV. We did not see any significant changes on the number of HFB in FAM001_DRAV or NBD in PAT001_DRAV, indicating that excitatory networks alone could not predict drug efficacy in DS patients (Fig. 5A). However, Valproic acid significantly decreased the number of HFB in FAM001_GEFS, while Carbamazepine significantly increased the number of HFB, in line with the clinical history. Indeed, Carbamazepine also aggravated the HFB phenotype in PAT001_GEFS, while the neuronal network remained unresponsive to Valproic acid, again in line with the clinical history of this patient (Fig. 5A, B). While PAT001_GEFS seizures were responsive to Topiramate, Topiramate aggravated the neuronal network phenotype. This could be explained by the fact that Topiramate most likely works by inhibiting Na^+^ -channels in excitatory cultures, and not by modulating GABA_A_ receptor-mediated currents. Although we could not uncover differences between GEFS+ and DS in the basal phenotypes, the neuronal network remained unresponsive to ASM in DS patients, while drug response in GEFS+ patients could be predicted in a clinical relevant matter, further implicating excitatory neurons as a phenotypic contributor.

**Figure 5.**
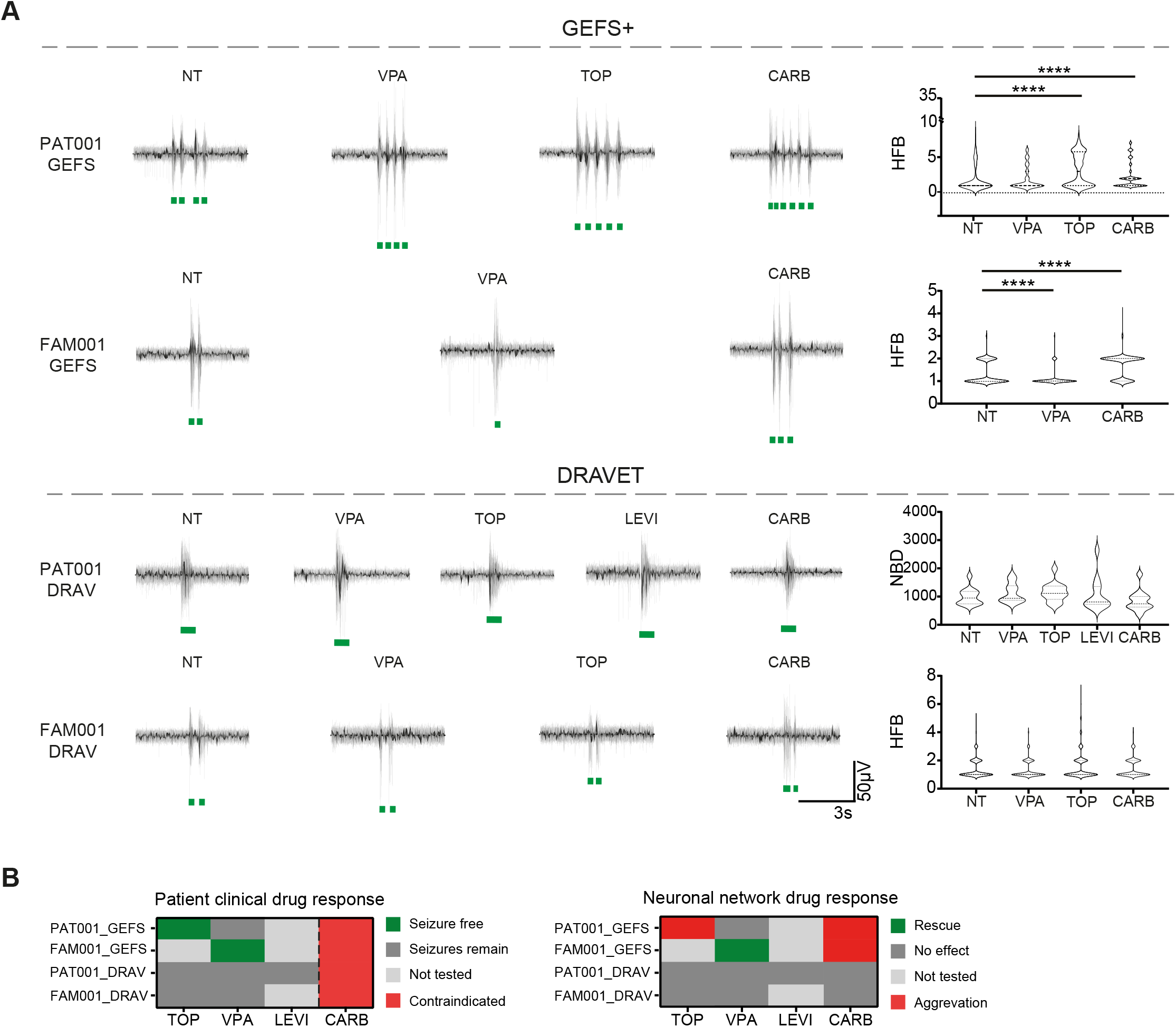
GEFS+, but not DS-derived neuronal networks respond to ASM. A) Representative burst traces of GEFS+ patients (top) and DS patients (bottom) including quantification of number of high frequency bursts (HFB) for PAT001_GEFS, FAM001_GEFS and FAM001_DRAV lines, and network burst duration (NBD) for PAT001_DRAV in non-treated (NT), or treated with 10 μM Topiramate (TOP), Valproic acid (VPA), Levetiracetam (LEVI), or Carbamazepine (CARB) conditions. HFB quantification for PAT001_DRAV was not included, since this line does not show HFB. Dashed line represents median, dotted line represents quartiles. B) Visualization of the clinical anti-seizure medication (ASM) response of each included patient (left), where red represents general contra-indication, light grey represents ASM not tested in this patient, dark grey represents the ASM that reduces seizure frequency, although the patient was not completely seizure free, and green represents the ASM with which the patient was completely seizure free; and visualization of the response of the neuronal networks to the same ASM (right), where red represents aggravation of the network phenotype, light grey represents ASM not tested in this patient neuronal network, dark gray indicates no change in the network phenotype, and green represents the ASM that (partially) rescues the network phenotype. N = number of recordings/batches, PAT001_GEFS_NT_ n = 13/5, PAT001_GEFS_VAL_ n = 13/5, PAT001_GEFS_TOP_ n = 13/5, PAT001_GEFS_CARB_ n = 13/5, PAT001_DRAV_NT_ n = 13/5, PAT001_DRAV_VAL_ n = 15/5, PAT001_DRAV_TOP_ n = 14/6, PAT001_DRAV_LEVI_ n = 12/5, PAT001_DRAV_CARB_ n = 13/4, FAM001_GEFS_NT_ n = 18/6, FAM001_GEFS_VAL_ n = 24/6, FAM001_GEFS_CARB_ n = 19/4, FAM001_DRAV_NT_ n = 7/3, FAM001_DRAV_TOP_ n = 9/3, FAM001_DRAV_VAL_ n = 10/3 FAM001_DRAV_CARB_ n = 6/1. Significance calculated with ANOVA with Kruskal-Wallis test with Dunss correction for multiple comparisons. \**P* = 0.05, \*\**P* = 0.01, \*\*\**P* = 0.001, \*\*\*\**P*<0.0001. All means, SEM and test statistics are listed in table 1.

## Discussion

The contribution of excitatory neurons to the DS phenotype has been heavily debated. We wondered whether the observed excitatory phenotype in some studies, but not in others, might be attributed to differences in patient clinical phenotype or the type of mutation. Here, we provide further evidence for the involvement of excitatory neurons in DS, in line with previous reports^20,26,27^, in four independent ways. First, we uncovered mutation-specific network phenotypes, while we could not distinguish patient clinical backgrounds. Specifically, the neuronal network phenotype did not differ between patients from one family with the same *SCN1A* mutation, but with different clinical phenotypes. Rather, the neuronal network phenotype from this family clustered together with that of an independent patient line, with a similar missense mutation. Indeed, mutations in the voltage sensing domain were distinguished from mutations in the pore domain. Second, we showed that the patient-derived network phenotypes correspond to the response of naïve networks to pro convulsive compounds. Third, we uncovered a previously unidentified excitatory neuronal network phenotype, HFB, which was aggravated by DS clinical-relevant trigger febrile temperatures. Finally, that GEFS+ patient-derived neuronal networks responded to ASM Valproic acid and Carbamazepine in a clinically relevant manner further supported the implication of excitatory neurons in DS.

While mutation specific changes have not been previously described in *SCN1A*-deficient excitatory neuronal networks, previous research uncovered variant specific alterations in Na_V_1.1 properties on a single cell level, in both excitatory and inhibitory neurons^38–41^. The type or position of the mutation in the channel can predict if the mutation leads to GoF or LoF ^8^. On a single cell level, we observed a significant decrease in Na^+^ -current density in DS-derived neurons with a missense mutation in the pore domain, matching the expected LoF properties of a pore-domain mutation^8^. However, we could not detect significant changes in Na^+^ -currents in neurons with mutations in the voltage sensing domain. We envisage two explanations. First, the elicited Na^+^ -currents were affected by space clamp artifacts^27^, caused by the expression of Na^+^ -channels at sites electrically distant from the neuronal soma, making it difficult to achieve complete space clamp. This is in line with numerous other studies investigating Na^+^ -currents in hiPSC-derived neurons^21–23,25–27^. Second, the expression of Na_V_1.1 in excitatory neurons might be insufficient to properly detect more subtle changes in Na_V_1.1 - currents, which could be overshadowed by other Na^+^ -channels working in concert. Small changes in Na_v_1.1 -currents might have a cumulative effect, explaining why we observed changes in activity on a neuronal network level. In particular, homozygous null *Scn1a^-/-^* mice, showed specific upregulation of Na_V_1.3 in hippocampal interneurons^13^. Finally, Na^+^ -channels can function as a dimer and display coupled gating. Interaction with WT channels could therefore also influence channel kinetics^42^. Taken together, true mechanistic insights into the biophysical properties of each variant should be carefully measured in expression systems with proper voltage control.

Interestingly, we here reported that *SCN1A*-deficiency, and reduced Na^+^ -current density did not result in reduced firing, but rather increased excitability in *SCN1A^+/-^* -deficient neurons. This could be due to aforementioned upregulation of other Na_V_ -channels, but our data also point to a compensatory mechanism from K_V_ -channels. We uncovered significant changes in action potential decay kinetics and AHP in *SCN1A^+/-^* -deficient neurons. In addition, we observed a hyperpolarizing shift in ΔAHP in all patient-derived neurons that displayed HFB, suggesting similar underlying mechanisms. Moreover, blocking K_V_ -channels by 4-ap and Linopirdine resulted in a similar hyperactive neuronal network phenotype with HFB. Compensatory rebalancing in excitatory neurons was observed in other works, for example in a *Scn2a* mutant mouse model, the K_V_ -channel K_V_8.2 has shown to function as a genetic modifier, resulting in increased susceptibility to epilepsy in pyramidal cells^43^. Moreover, in another *Scn2a* mouse model, neocortical pyramidal cell hyperexcitability was explained by a reduction in hyperpolarizing K^+^ -currents^44^. Finally, in the *Scn1b^-/-^* DS mouse model, increased neuronal input resistance caused pyramidal cell hyperexcitability. This phenotype was rescued by ASM Retigabine, a K_V_ -channel opener that reduces neuronal input resistance^45^. To conclude, compensatory re-balancing of K_V_ -channels might explain why *SCN1A^+/-^* -deficient excitatory neurons appear as paradoxically hyperactive.

Both this study and previous works fundamentally indicate that *SCN1A* mutations do not exclusively affect inhibitory neurons. However, the role of inhibitory neurons in the pathophysiology of DS is indisputable, especially considering the broad group of ASM prescribed for DS that work on the GABAergic system. Nevertheless, we were interested if typical ASM could also affect the excitatory network phenotype, and if excitatory networks alone could already give an indicative prediction of ASM non-responders versus responders. We uncovered that GEFS+-patient derived neuronal networks, but not DS networks responded to ASM in a clinically relevant matter. It is essential to note that on a clinical level the DS patients in our cohort are not seizure free with ASM, while the GEFS+ patients are. It might be that an excitatory-only neuronal network is not the ideal model to predict more subtle changes in seizure frequency upon ASM administration. Future research should focus on the incorporation of inhibitory neurons to the network^46^, which might serve as a more ideal model for patient-specific drug response in the context of DS. That we were nonetheless able to distinguish GEFS+ and DS patients responses to ASM further supports the role of excitatory neurons to the phenotype. Through which mechanisms the ASM affects excitatory neurons is still unclear. The working mechanisms of Valproic acid are poorly understood and it predominantly functions through regional changes in GABA-concentration. Previous research uncovered that Valproic acid can also regulate pERK trough attenuation of the PKA system^47^. Interestingly, proteomic signature analysis of *Scn1a*^+/-^ mice revealed altered protein regulation in glutamatergic synapses specifically, implicating a dysregulation of the PKA pathway, before the seizure threshold^48^. It could be that Valproic acid alters excitatory neuronal networks through a similar mechanism. Moreover, Topiramate acts as a positive allosteric modulator of GABA_A_ receptor-mediated currents, but also inhibits Na_V_ -channels. In our excitatory only networks, Topiramate will clearly not affect GABAa mediated currents. Therefore, the sole target of Topiramate in our cultures are Na_V_ -channels, which is a contra indication for DS. This could explain the aggravated or unaffected excitatory network phenotype we observed.

To conclude, we here show that the implication of the excitatory neuronal network phenotype is mutation specific in DS, which is further supported by the response of the neuronal network to DS-clinically relevant features, and provide evidence that neuronal networks on MEAs can be leveraged for patient specific drug screening in the context of DS in the future.

## Data and material availability statement

All data and algorithms generated in this study can be requested from the corresponding author Nael Nadif Kasri. Aside from the hiPSC lines, this study did not generate unique new reagents. All reagents, vectors or cell lines used in this study are available from the corresponding author upon request with a completed Materials Transfer Agreement.

## Acknowledgements and funding information

This work was supported by the Netherlands Organisation for Health Research and Development ZonMw grant 91217055 (to H.v.B., N.N.K., M.M., J.V., and H.J.S.), the European Joint Programme on Rare Diseases JTC2020-SCN1A-up! (to N.N.K.), the Dutch epilepsiefonds WAR 18-02 (to N.N.K., J.V., M.M. and H.J.S).

## Short running title

Modeling Dravet syndrome

## Supplementary methods

### Patient clinical history

PAT001_DRAV was a female patient born after an uneventful pregnancy. She presented with status epilepticus at six months. Her seizures further developed into atonic seizures, tonic-clonic seizures, hemiclonic seizures (left side) and myoclonic seizures, including another status at nine months. Her seizures are currently treated by a combination therapy of Levetiracetam, Clobazam, Stiripentol and Valproic acid, but remain uncontrolled with periods of forceful disruptions. She was diagnosed with ADHD including behavioral problems and the Wechsler Intelligence Scale for Children (WISC-V) test determined mild intellectual disability (TIQ 69), including a disharmonic profile, low socio-emotional level, reduced attention span and quick sensory overstimulation. The patient further presented with difficulties in feeding and frequent urinary tract infections. Direct target sequencing for *SCN1A* mutations revealed a missense mutation in the pore domain at position c.4168G>A (p. Val1390Met) (DIII S5-S6) (Fig. 3B).

PAT001_GEFS was a female patient, and had her fist tonic-clonic seizure at 1 year, which further developed into bilateral tonic-clonic seizures. Her development was normal before her first seizure. WISC-III-NL testing determined low-average intelligence, and showed characteristics of ADHD. Her seizures were unresponsive to Valproic acid and Clobazam, but were completely absent with the administration of Topiramate. Her last seizure was triggered by a failed attempt to phase out the Topiramate treatment. Family diagnostic screening revealed familial inheritance of a *SCN1A* missense mutation from the paternal line in the voltage sensing domain at position c.2576G>A (p.Arg859His) (DII S4) (Fig. 3B).

FAM001_DRAV and FAM001_GEFS belonged to the same family, and inherited a missense mutation in the voltage sensing domain of *SCN1A* at the c.3926T>G (p. Leu1309Arg) (DIIIS4) position from their father (FAM001_FS) (Fig. 3B), who only had febrile seizures as a child and no further comorbidities.

FAM001_DRAV was a male patient and born after emergency caesarian section after a normal pregnancy. His first seizure was at six months, and further developed into tonic-clonic and myoclonic seizures. The patient responded well to Topiramate and Valproic acid, and was seizure free for 5 years between the age of 11 and 16. The seizures resumed with tonic-clonic seizures, but were low in frequency, often triggered by physical activity. His development was normal before the first seizure. The patient further presented with mild intellectual disability, determined by WISC-III-NL testing (VIQ 67-PIQ 64). Moreover, the patient was diagnosed with behavioral problems, autism and ADHD.

FAM001_GEFS was a female patient, born after an uneventful pregnancy. Her seizures debuted at 15 months of age, and developed into tonic-clonic seizures and focal seizures that progressed into generalized seizures with a duration of 20 minutes. The seizures responded well to Valproic acid, and she was seizure free at the age of 4 years. Valproic acid was discontinued, and she remained seizure free. The patient further had low-average intelligence, was remarkably slow in processing and thinking, and was easily distracted, for which she received methylphenidate between the ages 7 and 12.

### Neuronal differentiation

24-well plates containing acid treated coverslips or Multi-Electrode Arrays (MEAs) were coated with Poly-L-ornithine (50 μg/ml, Sigma-aldrich, #P3655) for 3 hours at 37 °C/5%CO^2^, followed by three washes with mili-Q and an overnight coating with Biolaminin (5 μg/ml) at 4 °C. At DIV0, hiPSCs were washed with PBS, followed by a 3-5 minutes incubation with Accutase (Sigma-Aldrich, #A6964) at 37 °C/5%CO^2^. HiPSCs were centrifuged in DMEM/F-12 (Gibco, #11320074) for 5 min at 90x G, and resuspended until single cells in E8 medium containing doxycycline (4 μg/ml, Sigma-Aldrich, #D2975000) and RevitaCell (10 μg/ml). Cells were plated at a final density of 20.000 cells per well. At DIV 1, the medium was exchanged for DMEM/F12 supplemented with N-2 (1%, Gibco, #17502048), MEM Non-essential amino acids (1%, Sigma-Aldrich, #M7145), NT3 (10 ng/ml, Promokine #C-66425), recombinant human brain-derived neurotrophic factor (BDNF) (10 ng/ml, Promokine, # C-6621), DAPT (10 μM, Sigma-Aldrich, #565770), doxycycline (4 μg/ml) and mouse laminin (0.2 μg/ml, Sigma, #L2020). At DIV 2, E18 rat astrocytes were added in a 1:1 ratio. At DIV 3, the medium was replaced with Neurobasal medium (Gibco, #21103049), supplemented with B-27 (20 μg/ml, Thermo Fisher, #17504044), Glutamax (10 μg/ml Thermo Fisher, #5050061), primocin (0.1 μg/ml), NT3 (10 ng/ml), BDNF (10 ng/ml), doxycycline (4 μg/ml) and Cytosine β-D-arabinofuranoside (2 μM, Sigma-Aldrich, #C176) to remove proliferating cells. Over the following days, 50% of the medium was replaced every 2 days. From DIV6 onwards the medium was supplemented with NT3, BDNF, and doxycycline, which from DIV10 onwards was supplemented with FBS (2,5%, Sigma, #F7524) to enhance astrocyte viability. From DIV13 onwards, doxycycline was removed. This final composition was used for the remainder of the differentiation culture, which lasted up to DIV51. Throughout culturing, the neuronal cultures were kept at 37°C/5% CO^2^.

---

**Figure S1.**
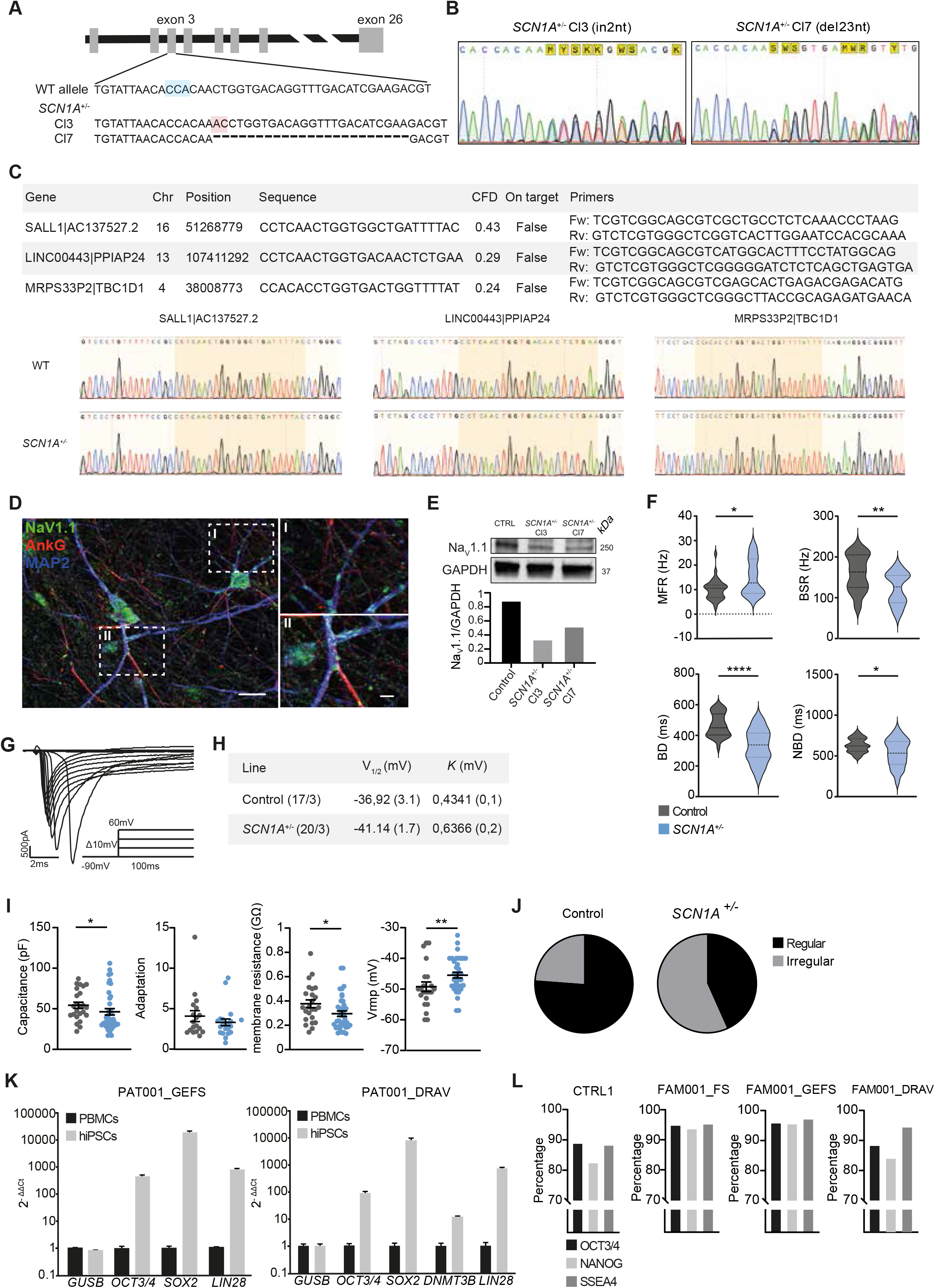
Description of Na_V_1.1 expression and Na^+^ -current dynamics in *SCN1A^+/-^* -deficient neurons. A) Schematic overview of generated *SCN1A^+/-^* deficient lines. Blue represents PAM site, red represents insertion. B) Chromatograms of sequencing results from *SCN1A^+/-^* Cl3 (insertion of 2 nucleotides) and *SCN1A^+/-^* Cl7 (deletion of 23 nucleotides) C) Top 3 potential off-target sites have been sequenced and no mutations were detected. D) Representative image of Na_V_1.1 expression in control day *in vitro* (DIV) 49 neurons; dendritic marker MAP2 in blue, axon initial segment marker AnkG in red and Na_V_1.1 in green. Scale bar represents 20 μm, and 40 μm (inset) E) Quantification of Na_V_1.1 protein levels relative to GAPDH in DIV49 control and *SCN1A^+/-^* -deficient neurons (cropped image) n = 2. F) Quantification of network parameters including: mean firing rate (MFR), burst spike rate (BSR), and (network) burst duration ((N)BD). n = number of recordings/batches: control n = 33/5 and *SCN1A^+/-^* n = 23/3, Mann-Withney test. \**P* = 0.05, \*\**P* = 0.01, \*\*\**P* = 0.001, \*\*\*\**P*<0.0001. All means, SEM and test statistics are listed in table 1. G) Representative Na^+^ -current traces and stimulation paradigm (inset): stepwise protocol from a −90 mV holding potential to a maximum test-pulse of 60 mV in increments of 10 mV. H) Table describing the half maximum of activation (V_1/2_) and slope (*k*) after Bolzmann fitting. I) Analysis of active and passive membrane properties including: membrane capacitance, spike frequency adaptation, defined as the inter spike interval (ISI) between the second and third action potential, divided by the ISI of the second half of the sweep, membrane resistance, and resting membrane potential (Vrmp). J) quantification of regular vs irregular firing patterns in control and *SCN1A^+/-^* neurons. Irregularity was defined as any trace with two action potentials with a longer inter spike interval (ISI) than predicted based on previous and next ISIs in the same trace. K) Pluripotency marker expression in hiPSCs relative to PBMC (peripheral mononuclear blood cells) determined using Quantitative real time PCR in PAT001_DRAV and PAT001_GEFS lines. Delta ct levels of octamer-binding transcription factor 3/4 (*OCT3/4)*, SRY-box 2 (*SOX2*), *DNMT3B*, and *LIN28*, using glucuronidase beta *GUSB* as housekeeping gene, displayed as the relative gene expression normalized to *GUSB*. L) Flow cytometry quantification of pluripotency markers in CTRL1, FAM001_FS, FAM001_GEFS and FAM001_DRAV lines. Data represented as percentage of marker positive nuclei.

**Figure S2.**
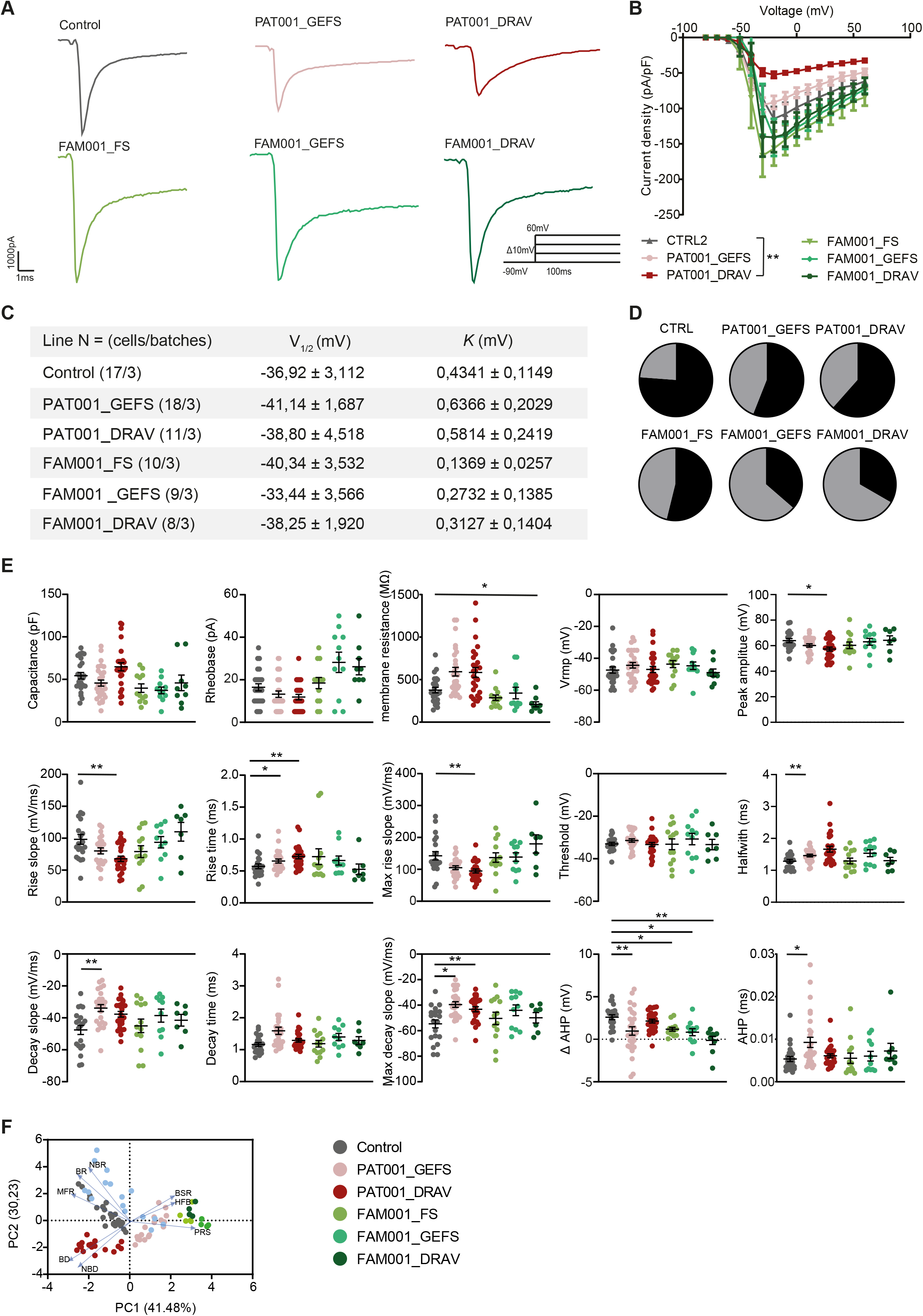
Quantification of Na^+^ -currents and action potential intrinsic properties in DS-patient derived neurons. A) Representative Na^+^ -current traces of control and patient-derived neurons at *day in vitro* (DIV) 49. Stimulation paradigm (inset): stepwise protocol from a −90 mV holding potential to a maximum test-pulse of 60 mV in increments of 10 mV B) Current-density plot of Na^+^ -current recordings from control and patient derived neurons. C) Boltzmann fitting of Na^+^ -currents measured in control and patient-derived neurons, describing the half maximum of activation (V_1/2_) and slope (*k*). D) Quantification of regular vs irregular firing patterns in control and patient-derived neurons. Irregularity was defined as any trace with two action potentials with a longer inter spike interval (ISI) than predicted based on previous and next ISIs in the same trace E) Quantification of action potential intrinsic properties, membrane capacitance, rheobase, membrane resistance, resting membrane potential (Vrmp), peak amplitude, rise slope, rise time, maximum rise slope, threshold, halfwidth, decay slope, decay time, max decay slope, after hyper polarization time (AHP), and ΔAHP, defined as the difference between the AHP of the first action potential and the second action potential in the same sweep. n = number of recordings/batches CTRL2 n = 20/3, PAT001_GEFS n = 25/3, PAT001_DRAV n = 26/3, FAM001_FS n = 13/2 FAM001_GEFS n = 11/2 FAM001_DRAV n = 7/2, ANOVA with Bonferroni correction for multiple comparisons. Data is shown as mean ±SEM. *p = 0.05, **p = 0.01, ***p = 0.001, ****P<0.0001. All means, SEM and test statistics are listed in table 1. F) PCA-plot of 8 MEA parameters, including mean firing rate (MFR), burst rate (BR), NBR, percentage of random spikes (PRS), (network) burst duration ((N)BD), high frequency bursts (HFB), burst spike rate (BSR), interburst interval (IBI) and network inter burst interval (NIBI), showing parameters that explain the differences in network behavior between control and patient lines during febrile temperatures. Blue arrows indicate loadings. N = number of recordings/batches control n = 20/3, PAT001_GEFS n = 12/2, PAT001_DRAV n = 22/3 FAM001_FS n = 5/1, FAM001_GEFS n = 9/2, FAM001_DRAV n = 8/2.

